## Supplementary Figures and Table for "Functional Implications of the Exon 9 Splice Insert in GluK1 Kainate Receptors"

### Figure supplements

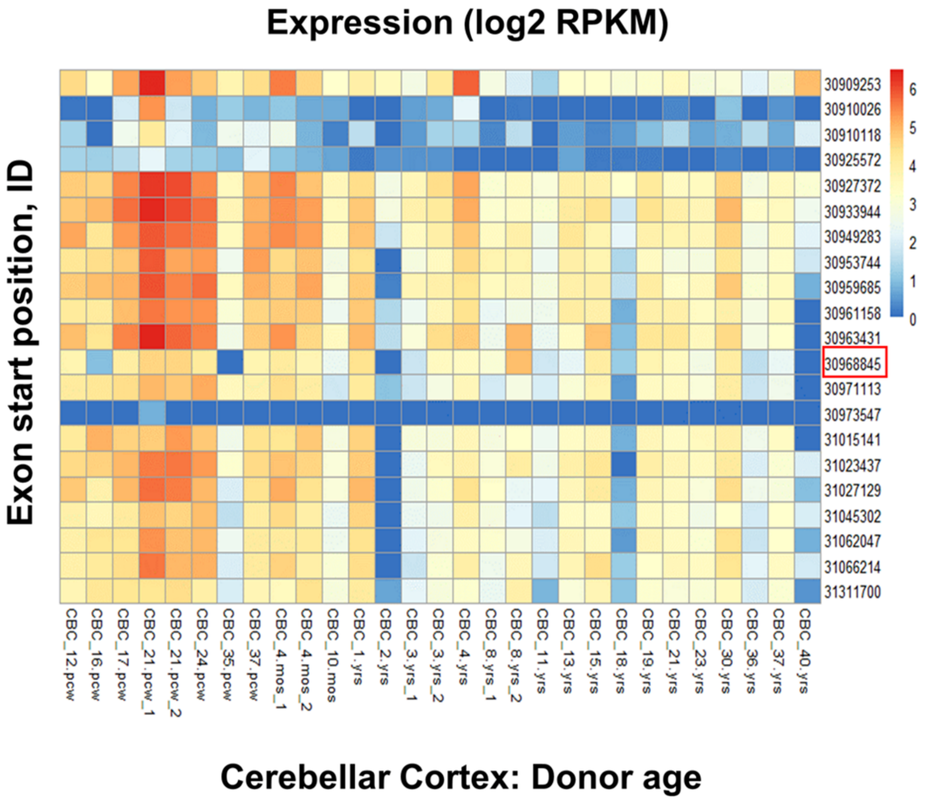


**Figure 1-figure supplement 1.** Representative RNA-Seq analysis (BrainSpan atlas) of the Cerebellar cortex (CBC) demonstrates the high expression of exon 9 in the human brain. To delve deeper into the high expression areas from **Figure 1**, cerebellar cortex was chosen as an example to understand the expression pattern of Exon 9 with respect to other GluK1 exons. The heat map displays abundance of the GluK1 exons at different stages of human life. Exon 9 (Exon start position: 30968845; marked by a red box) expression coincides with reported high expression of GluK1 kainate receptors in early developmental stages of humans. Blue and red color indicate zero and maximum expression respectively.

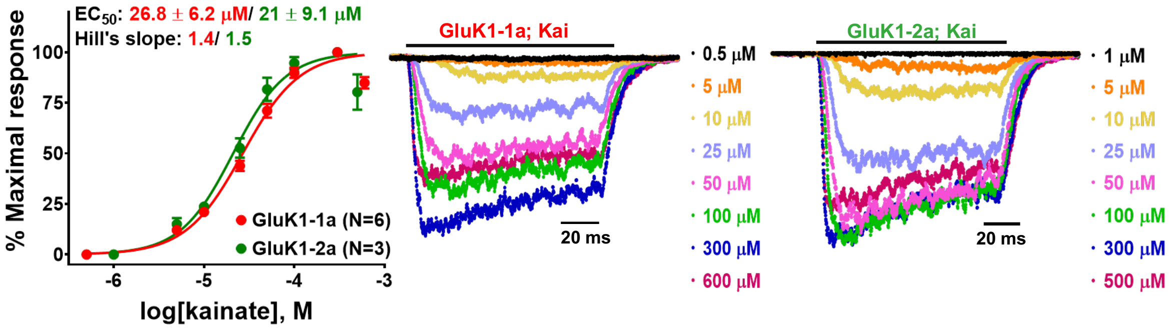


**Figure 2-figure supplement 1.** Kainate-evoked responses for GluK1 receptors. The figure demonstrates the normalized kainate dose-response curves for GluK1-1a and GluK1-2a along with their representative traces.

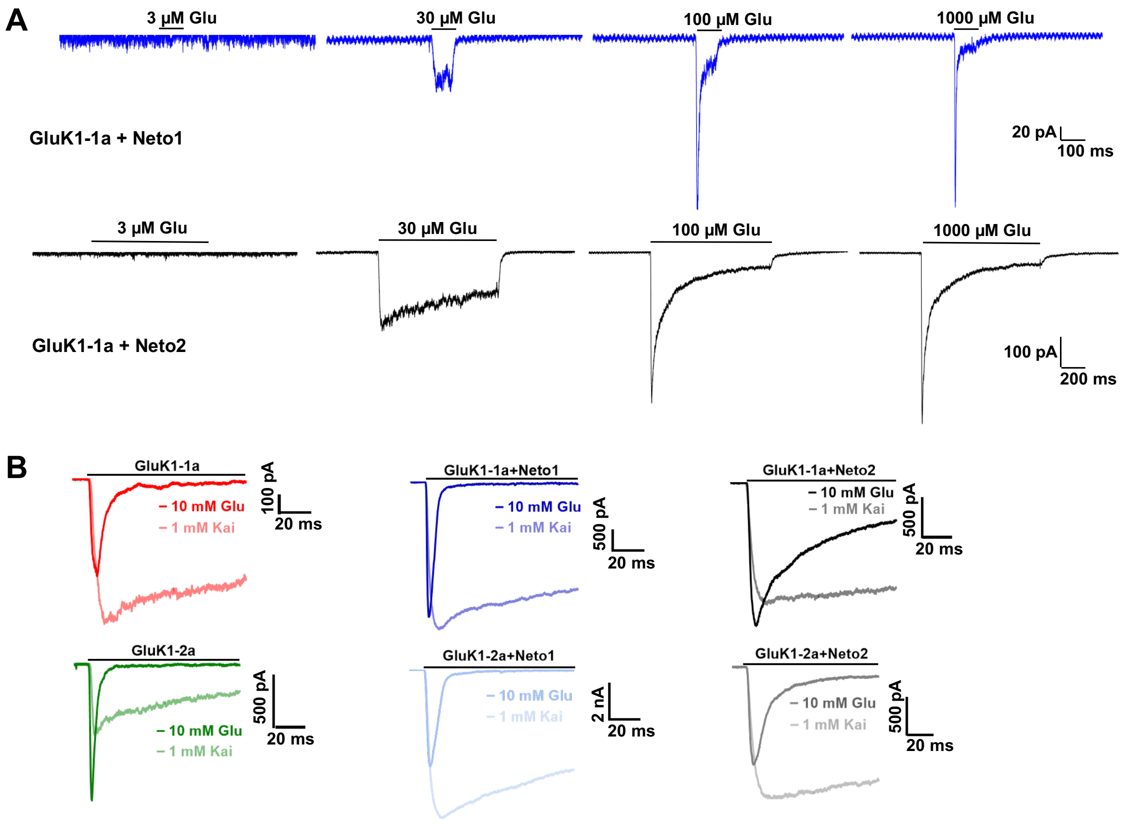


**Figure 3-figure supplement 1.** Effect of Neto proteins on agonist-evoked responses of GluK1 receptors. **(A)** Displays representative traces for GluK1-1a in the presence of Neto1 (blue) or Neto2 (black) with varying concentrations of glutamate (3 µM, 30 µM, 100 µM & 1000 µM). Ligand application for receptors with Neto1 and Neto2 was 100 ms, and 1000 ms, respectively. **(B)** Representative 100 ms traces for currents evoked in presence of glutamate vs. kainate in the absence and presence of Neto proteins.

| $K_{368}$**ASGEVSKHLYKVWK**$_{382}$ | | **Forward primer (5'-3')** | **Reverse primer (5'-3')** |
|---|---|---|---|
| **Common cloning primers (AfeI/BsiWI: RE sites)** | | CTGAAGCAGCGCTGATGTACG | CTTTCTCCCGTACGTATGTGATGG |
| **Ala mutations (Charge neutral mutations)** | | | |
| $K_{375}H_{376}$-A | KASGEVSAALYKVWK | GCCTCTGGTGAAGTGTCTGCCGCTTTGTATAAAGTGTGGAAG | CTTCCACACTTTATACAAAGCGGCAGACACTTCACCAGAGGC |
| $K_{375/379/382}$-A | KASGEVSAHLYAVWA | GCCTCTGGTGAAGTGTCTGCCCACTTGTATGCCGTGTGGGCCAAGATTGGGATTTGGAAC | GTTCCAAATCCCAATCTTGGCCCACACGGCATACAAGTGGGCAGACACTTCACCAGAGGC |
| $Y_{378}V_{380}W_{381}$-A | KASGEVSKHLAKAAK | GAAGTGTCTAAACACTTGGCTAAAGCGGCCAAGAAGATTGGGATTTGG | CCAAATCCCAATCTTCTTGGCCGCTTTAGCCAAGTGTTTAGACACTTC |
| $K_{375}$HLYKVWK$_{382}$-8A | KASGEVSAAAAAAAA | GCCTCTGGTGAAGTGTCTGCCGCTGCAGCTGCAGCCGCAGCTAAGATTGGGATTTGGAAC | GTTCCAAATCCCAATCTTAGCTGCGGCTGCAGCTGCAGCGGCAGACACTTCACCAGAGGC |
| **Glu mutations (Charge reversal mutations)** | | | |
| $K_{375}H_{376}$-E | KASGEVSEELYKVWK | GCCTCTGGTGAAGTGTCTGAGGAATTGTATAAAGTGTGGAAG | CTTCCACACTTTATACAATTCCTCAGACACTTCACCAGAGGC |
| $H_{376}$-E | KASGEVSKELYKVWK | GCCTCTGGTGAAGTGTCTAAAGAATTGTATAAAGTGTGGAAG | CTTCCACACTTTATACAATTCTTTAGACACTTCACCAGAGGC |
| $K_{375/379/382}H_{376}$-E | KASGEVSEELYEVWE | GCCTCTGGTGAAGTGTCTGAGGAATTGTATGAGGTGTGGGAGAAGATTGGGATTTGGAAC | GTTCCAAATCCCAATCTTCTCCCACACCTCATACAATTCCTCAGACACTTCACCAGAGGC |
| $K_{368/375/379/382}H_{376}$-E | EASGEVSEELYEVWE | AAAGAGGAAGGAACTGAAGAGGCCTCTGGTGAAGTGTCT | AGACACTTCACCAGAGGCCTCTTCAGTTCCTTCCTCTTT |
| $K_{368}$-E | EASGEVSKKLYKVWK | AAAGAGGAAGGAACTGAAGAGGCCTCTGGTGAAGTGTCT | AGACACTTCACCAGAGGCCTCTTCAGTTCCTTCCTCTTT |

**Figure 4-table supplement 1.** GluK1-1a ATD splice mutants. The various mutants used in the study and the primer sequences to make charge-neutral and charge-reversal mutants in GluK1-1a are tabulated.

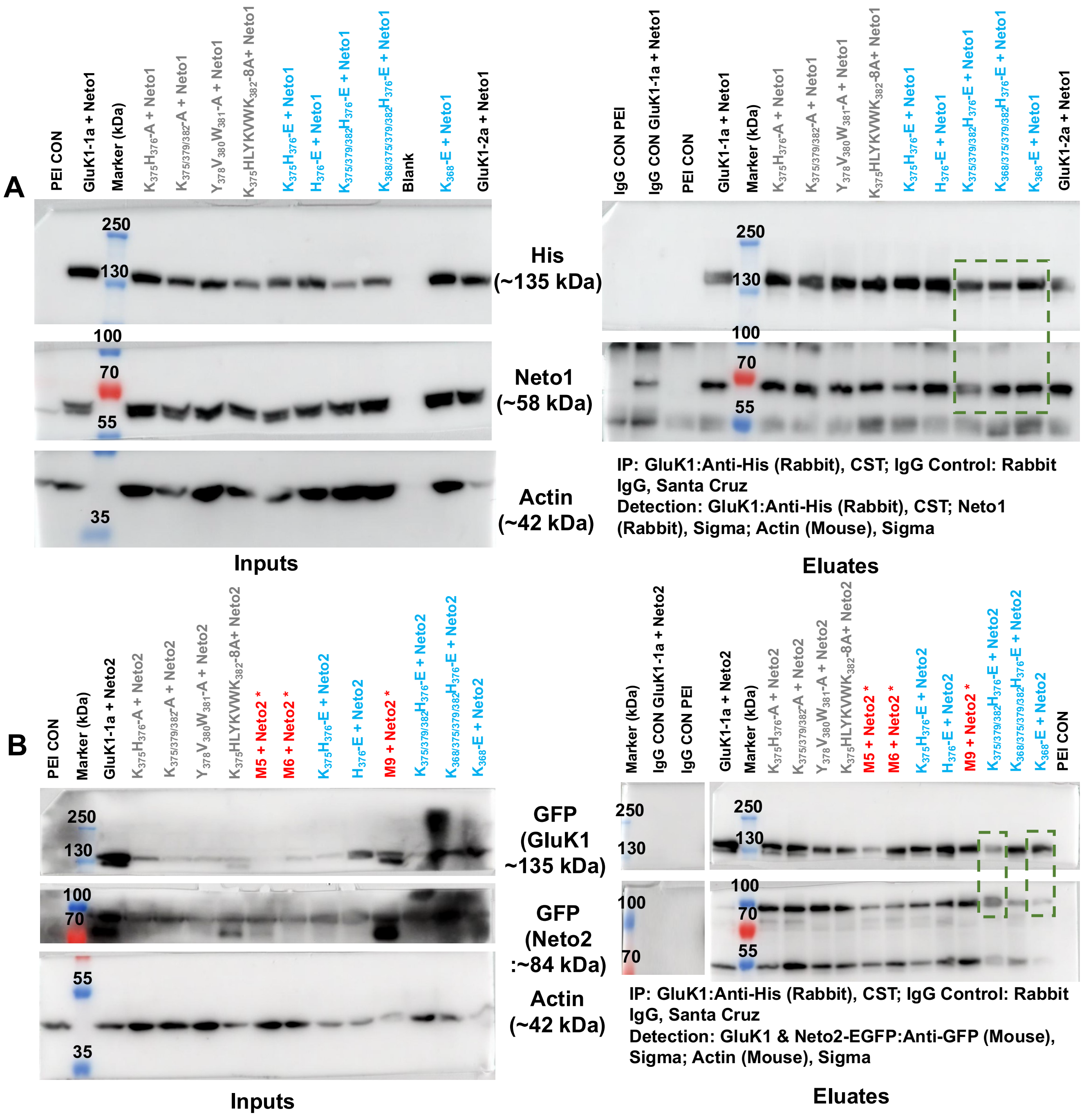


**Figure 5-figure supplement 1.** Co-immunoprecipitation analysis of GluK1-1a splice mutants and Neto proteins. **(A & B)** represent the raw western blots for receptor pull-down experiments using His-antibody (CST). The left and right panels show the inputs and eluates for the co-IP.

Actin was used as an internal control. The antibodies used to detect receptor or Neto proteins have been indicated. Rabbit IgG controls were set up using WT receptor with Neto1 or Neto2 and negative control along with the test samples to check for non-specific interactions. All the experiments were done in triplicates. Charge-neutral (Ala) mutants are labeled in grey, charge reversal (Glu) mutants in cyan and marker, and controls in black. Red asterisks are mutants not used for further studies. The molecular weight of markers and the expected proteins has been indicated. Green boxes display that the mutant receptors shown in functional analysis were able to interact with both Neto proteins efficiently when co-transfected with Neto1 or Neto2.

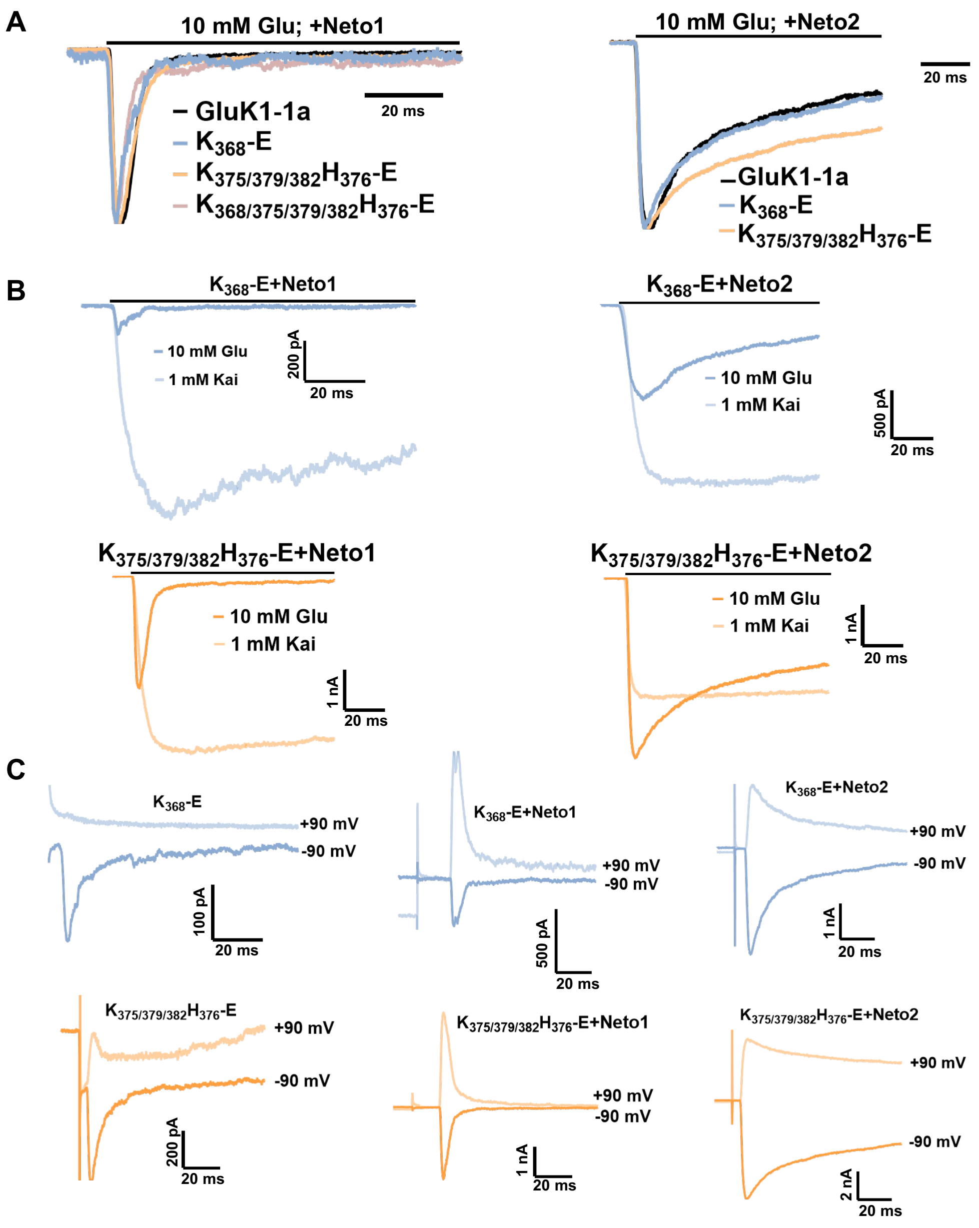


**Figure 5-figure supplement 2.** Representative traces for the GluK1 mutant receptors with Neto proteins. **(A)** Demonstrates the normalized traces for wild-type and mutant receptors with Neto1/2 for glutamate evoked desensitization. **(B)** Displays representative traces for glutamate vs. kainate evoked responses in the presence of Neto1 or Neto2. **(C)** Representative 100 ms traces for currents evoked at positive (+90 mV) and negative (-90 mV) potentials in the absence and presence of Neto proteins.

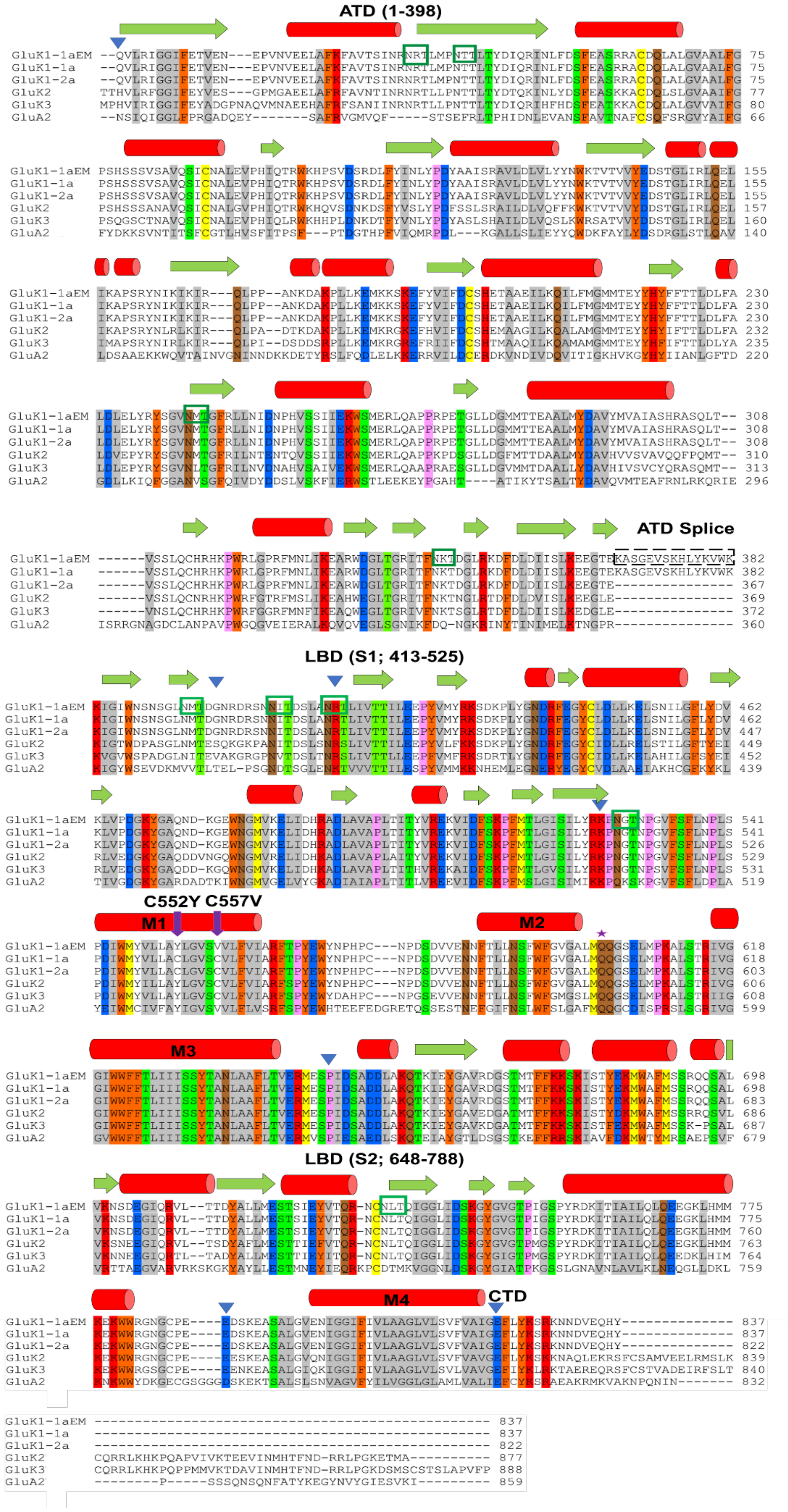

ATD (1-398)
GluK1-1aEM --QVLRIGGIFETVEN---EPVNVEELAFKFAVTSINRNRTLMPNTTLTYDIQRINLFDSFEASRRACDQLALGVAALFG 75
GluK1-1a --QVLRIGGIFETVEN---EPVNVEELAFKFAVTSINRNRTLMPNTTLTYDIQRINLFDSFEASRRACDQLALGVAALFG 75
GluK1-2a --QVLRIGGIFETVEN---EPVNVEELAFKFAVTSINRNRTLMPNTTLTYDIQRINLFDSFEASRRACDQLALGVAALFG 75
GluK2 TTHVLRFGGIFEYVES---GPMGAEELAFRFAVNTINRNRTLLPNTTLTYDTQKINLYDSFEASKKACDQLSLGVAAIFG 77
GluK3 MPHVIRIGGIFEYADGPNAQVMNAEEHAFRFSANIINRNRTLLPNTTLTYDIQRIHFHDSFEATKKACDQLALGVVAIFG 80
GluA2 --NSIQIGGLFPRGADQEY-------SAFRVGMVQF-----STSEFRLTPHIDNLEVANSFAVTNAFCSQFSRGVYAIFG 66
GluK1-1aEM PSHSSSVSAVQSICNALEVPHIQTRWKHPSVDSRDLFYINLYPDYAAISRAVLDLVLYYNWKTVTVVYEDSTGLIRLQEL 155
GluK1-1a PSHSSSVSAVQSICNALEVPHIQTRWKHPSVDSRDLFYINLYPDYAAISRAVLDLVLYYNWKTVTVVYEDSTGLIRLQEL 155
GluK1-2a PSHSSSVSAVQSICNALEVPHIQTRWKHPSVDSRDLFYINLYPDYAAISRAVLDLVLYYNWKTVTVVYEDSTGLIRLQEL 155
GluK2 PSHSSANAVQSICNALGVPHIQTRWKHQVSDNKDSFYVSLYPDFSSLSRAILDLVQFFKWKTVTVVYDDSTGLIRLQEL 157
GluK3 PSQGSCTNAVQSICNALEVPHIQLRWKHHPLDNKDTFYVNLYPDYASLSHAILDLVQSLKWRSATVVYDDSTGLIRLQEL 160
GluA2 FYDKKSVNTITSFCGTLHVSFITPSF---PTDGTHPFVIQMRPDL---KGALLSLIEYYQWDKFAYLYDSDRGLSTLQAV 140
GluK1-1aEM IKAPSRYNIKIKIR---QLPP--ANKDAKPLLKEMKKSKEFYVIFDCSHETAAEILKQILFMGMMTEYYHYFFTTLDLFA 230
GluK1-1a IKAPSRYNIKIKIR---QLPP--ANKDAKPLLKEMKKSKEFYVIFDCSHETAAEILKQILFMGMMTEYYHYFFTTLDLFA 230
GluK1-2a IKAPSRYNIKIKIR---QLPP--ANKDAKPLLKEMKKSKEFYVIFDCSHETAAEILKQILFMGMMTEYYHYFFTTLDLFA 230
GluK2 IKAPSRYNLRLKIR---QLPA--DTKDAKPLLKEMKRGKEFHVIFDCSHEMAAGILKQALAMGMMTEYYHYIFTTLDLFA 232
GluK3 IMAPSRYNIRLKIR---QLPI--DSDDSRPLLKEMKRGREFRIIFDCSHTMAAQILKQAMAMGMMTEYYHFIFTTLDLYA 235
GluA2 LDSAAEKKWQVTAINVGNINNDKKDETYRSLFQDLELKKERRVILDCERDKVNDIVDQVITIGKHVKGYHYIIANLGFTD 220
GluK1-1aEM LDLELYRYSGVNMTGFRLLNIDNPHVSSIIEKWSMERLQAPPRPETGLLDGMMTTEAALMYDAVYMVAIASHRASQLT-- 308
GluK1-1a LDLELYRYSGVNMTGFRLLNIDNPHVSSIIEKWSMERLQAPPRPETGLLDGMMTTEAALMYDAVYMVAIASHRASQLT-- 308
GluK1-2a LDLELYRYSGVNMTGFRLLNIDNPHVSSIIEKWSMERLQAPPRPETGLLDGMMTTEAALMYDAVYMVAIASHRASQLT-- 308
GluK2 LDVEPYRYSGVNMTGFRILNTENTQVSSIIEKWSMERLQAPPKPDSGLLDGFMTTDAALMYDAVHVVSVAVQQFPQMT-- 310
GluK3 LDLEPYRYSGVNLTGFRILNVDNAHVSAIVEKWSMERLQAAPRAESGLLDGVMMTDAALLYDAVHIVSVCYQRASQMT-- 313
GluA2 GDLLKIQFGGANVSGFQIVDYDDSLVSKFIERWSTLEEKEYPGAHT----ATIKYTSALTYDAVQVMTEAFRNLRKQRIE 296
ATD Splice
GluK1-1aEM ------VSSLQCHRHKPWRLGPRFMNLIKEARWDGLTGRITFNKTDGLRKDFDLDIISLKEEGTEKASGEVSKHLYKVWK 382
GluK1-1a ------VSSLQCHRHKPWRLGPRFMNLIKEARWDGLTGRITFNKTDGLRKDFDLDIISLKEEGTEKASGEVSKHLYKVWK 382
GluK1-2a ------VSSLQCHRHKPWRLGPRFMNLIKEARWDGLTGRITFNKTDGLRKDFDLDIISLKEEGTE--------------- 367
GluK2 ------VSSLQCNRHKPWRFGTRFMSLIKEAHWEGLTGRITFNKTNGLRTDFDLDVISLKEEGLE--------------- 369
GluK3 ------VNSLQCHRHKPWRFGGRFMNFIKEAQWEGLTGRIVFNKTSGLRTDFDLDIISLKEDGLE--------------- 372
GluA2 ISRRGNAGDCLANPAVPWGQGVEIERALKQVQVEGLSGNIKFDQ-NGKRINYTINIMELKTNGPR--------------- 360
LBD (S1; 413-525)
GluK1-1aEM KIGIWNSNSGLNMTDGNRDRSNNITDSLANRTLIVTTILEEPYVMYRKSDKPLYGNDRFEGYCLDLLKELSNILGFLYDV 462
GluK1-1a KIGIWNSNSGLNMTDGNRDRSNNITDSLANRTLIVTTILEEPYVMYRKSDKPLYGNDRFEGYCLDLLKELSNILGFLYDV 462
GluK1-2a KIGIWNSNSGLNMTDGNRDRSNNITDSLANRTLIVTTILEEPYVMYRKSDKPLYGNDRFEGYCLDLLKELSNILGFLYDV 447
GluK2 KIGTWDPASGLNMTESQKGKPANITDSLSNRSLIVTTILEEPYVLFKKSDKPLYGNDRFEGYCIDLLRELSTILGFTYEI 449
GluK3 KVGVWSPADGLNITEVAKGRGPNVTDSLTNRSLIVTTLLEEPFVMFRKSDRTLYGNDRFEGYCIDLLKELAHILGFSYEI 452
GluA2 KIGYWSEVDKMVVTLTEL-PSGNDTSGLENKTVVVTTILESPYVMMKKNHEMLEGNERYEGYCVDLAAEIAKHCGFKYKL 439
GluK1-1aEM KLVPDGKYGAQND-KGEWNGMVKELIDHRADLAVAPLTITYVREKVIDFSKPFMTLGISILYRKPNGTNPGVFSFLNPLS 541
GluK1-1a KLVPDGKYGAQND-KGEWNGMVKELIDHRADLAVAPLTITYVREKVIDFSKPFMTLGISILYRKPNGTNPGVFSFLNPLS 541
GluK1-2a KLVPDGKYGAQND-KGEWNGMVKELIDHRADLAVAPLTITYVREKVIDFSKPFMTLGISILYRKPNGTNPGVFSFLNPLS 526
GluK2 RLVEDGKYGAQDDVNGQWNGMVRELIDHKADLAVAPLAITYVREKVIDFSKPFMTLGISILYRKPNGTNPGVFSFLNPLS 529
GluK3 RLVEDGKYGAQDD-KGQWNGMVKELIDHKADLAVAPLTITHVREKAIDFSKPFMTLGVSILYRKPNGTNPSVFSFLNPLS 531
GluA2 TIVGDGKYGARDADTKIWNGMVGELVYGKADIAIAPLTITLVREEVIDFSKPFMSLGISIMIKKPQKSKPGVFSFLDPLA 519
C552Y C557V
M1
M2
GluK1-1aEM PDIWMYVLLAYLGVSVVLFVIARFTPYEWYNPHPC---NPDSDVVENNFTLLNSFWFGVGALMQQGSELMPKALSTRIVG 618
GluK1-1a PDIWMYVLLACLGVSCVLFVIARFTPYEWYNPHPC---NPDSDVVENNFTLLNSFWFGVGALMQQGSELMPKALSTRIVG 618
GluK1-2a PDIWMYVLLACLGVSCVLFVIARFTPYEWYNPHPC---NPDSDVVENNFTLLNSFWFGVGALMQQGSELMPKALSTRIVG 603
GluK2 PDIWMYILLAYLGVSCVLFVIARFSPYEWYNPHPC---NPDSDVVENNFTLLNSFWFGVGALMQQGSELMPKALSTRIVG 606
GluK3 PDIWMYVLLAYLGVSCVLFVIARFSPYEWYDAHPC---NPGSEVVENNFTLLNSFWFGMGSLMQQGSELMPKALSTRIIG 608
GluA2 YEIWMCIVFAYIGVSVVLFLVSRFSPYEWHTEEFEDGRETQSSESTNEFGIFNSLWFSLGAFMQQGCDISPRSLSGRIVG 599
M3
GluK1-1aEM GIWWFFTLIIISSYTANLAAFLTVERMESPIDSADDLAKQTKIEYGAVRDGSTMTFFKKSKISTYEKMWAFMSSRQQSAL 698
GluK1-1a GIWWFFTLIIISSYTANLAAFLTVERMESPIDSADDLAKQTKIEYGAVRDGSTMTFFKKSKISTYEKMWAFMSSRQQSAL 698
GluK1-2a GIWWFFTLIIISSYTANLAAFLTVERMESPIDSADDLAKQTKIEYGAVRDGSTMTFFKKSKISTYEKMWAFMSSRQQSAL 683
GluK2 GIWWFFTLIIISSYTANLAAFLTVERMESPIDSADDLAKQTKIEYGAVEDGATMTFFKKSKISTYDKMWAFMSSRRQSVL 686
GluK3 GIWWFFTLIIISSYTANLAAFLTVERMESPIDSADDLAKQTKIEYGAVKDGATMTFFKKSKISTFEKMWAFMSSK-PSAL 687
GluA2 GVWWFFTLIIISSYTANLAAFLTVERMVSPIESAEDLSKQTEIAYGTLDSGSTKEFFRRSKIAVFDKMWTYMRSAEPSVF 679
LBD (S2; 648-788)
GluK1-1aEM VKNSDEGIQRVL--TTDYALLMESTSIEYVTQR-NCNLTQIGGLIDSKGYGVGTPIGSPYRDKITIAILQLQEEGKLHMM 775
GluK1-1a VKNSDEGIQRVL--TTDYALLMESTSIEYVTQR-NCNLTQIGGLIDSKGYGVGTPIGSPYRDKITIAILQLQEEGKLHMM 775
GluK1-2a VKNSDEGIQRVL--TTDYALLMESTSIEYVTQR-NCNLTQIGGLIDSKGYGVGTPIGSPYRDKITIAILQLQEEGKLHMM 760
GluK2 VKSNEEGIQRVL--TSDYAFLMESTTIEFVTQR-NCNLTQIGGLIDSKGYGVGTPMGSPYRDKITIAILQLQEEGKLHMM 763
GluK3 VKNNEEGIQRTL--TADYALLMESTTIEYITQR-NCNLTQIGGLIDSKGYGIGTPMGSPYRDKITIAILQLQEEDKLHIM 764
GluA2 VRTTAEGVARVRKSKGKYAYLLESTMNEYIEQRKPCDTMKVGGNLDSKGYGIATPKGSSLGNAVNLAVLKLNEQGLLDKL 759
M4
CTD
GluK1-1aEM KEKWWRGNGCPE----EDSKEASALGVENIGGIFIVLAAGLVLSVFVAIGEFLYKSRKNNDVEQHY-------------- 837
GluK1-1a KEKWWRGNGCPE----EDSKEASALGVENIGGIFIVLAAGLVLSVFVAIGEFLYKSRKNNDVEQHY-------------- 837
GluK1-2a KEKWWRGNGCPE----EDSKEASALGVENIGGIFIVLAAGLVLSVFVAIGEFLYKSRKNNDVEQHY-------------- 822
GluK2 KEKWWRGNGCPE----EESKEASALGVQNIGGIFIVLAAGLVLSVFVAVGEFLYKSKKNAQLEKRSFCSAMVEELRMSLK 839
GluK3 KEKWWRGSGCPE----EENKEASALGIQKIGGIFIVLAAGLVLSVLVAVGEFIYKLRKTAEREQRSFCSTVADEIRFSLT 840
GluA2 KNKWWYDKGECGSGGGDSKEKTSALSLSNVAGVFYILVGGLGLAMLVALIEFCYKSRAEAKRMKVAKNPQNIN------- 832
GluK1-1aEM ------------------------------------------------- 837
GluK1-1a ------------------------------------------------- 837
GluK1-2a ------------------------------------------------- 822
GluK2 CQRRLKHKPQAPVIVKTEEVINMHTFND-RRLPGKETMA---------- 877
GluK3 CQRRLKHKPQPPMMVKTDAVINMHTFND-RRLPGKDSMSCSTSLAPVFP 888
GluA2 --------P-----SSSQNSQNFATYKEGYNVYGIESVKI--------- 859

**Figure 6-figure supplement 1.** Sequence alignment and construct design of GluK1-1a$_{EM}$. The EM construct was aligned with mature polypeptide sequences of wild-type (WT) rat GluK1-1a, GluK1-2a, GluK2, GluK3, and GluA2. The color scheme grouping was based on the similarity of residues, with grey (G, A, V, L, I), orange (F, Y, W), yellow (C, M), green (S, T), red (K, R, H), blue (D, E), brown (N, Q) and pink (P). The approximate domain boundaries and the residue numbers have been marked with blue inverted triangles and written in parenthesis. Green boxes mark predicted N-linked glycosylation sites (NXT). Purple arrows show cysteine mutations in the EM construct. The predicted secondary structure is shown above the sequence, with red cylinders for α-helix and green arrows for β-strand.

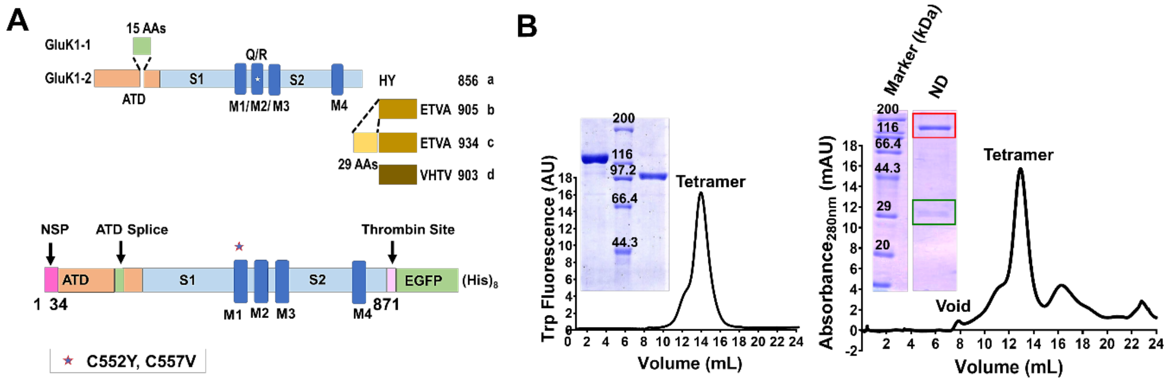


**Figure 6-figure supplement 2.** GluK1-1a$_{EM}$ construct design and purification. (**A**) Schematic representation of GluK1 splice variants present in the human brain along with a schematic for the GluK1-1a$_{EM}$ construct is shown. The N-terminal Signal Peptide (NSP; 1-34 residues), 15 amino acid splice insert in the Amino Terminal Domain (ATD), S1/S2 (Ligand Binding Domain, LBD), M1-M4 (Trans-Membrane Domain, TMD) and C-terminal thrombin site followed by EGFP and (His)$_8$ tags. The star in the M1 region denotes the point mutation of free cysteines at positions 552 and 557; the numbering of residues is based on the mature polypeptide. (**B**) Superose 6 size exclusion chromatography profiles for the purified protein in detergent micelles and nanodisc, respectively, are shown, and the position corresponding to void, receptor tetramer and empty nanodisc are indicated. SDS-PAGE gel images inset show the undigested and thrombin-digested protein for GluK1-1a$_{EM}$ in detergent and MSP1E3D1 (green box) co-eluting with GluK1-1a$_{EM}$ (red box), revealing a stable protein-nanodisc complex, respectively. ND indicates receptors in lipid nanodiscs.

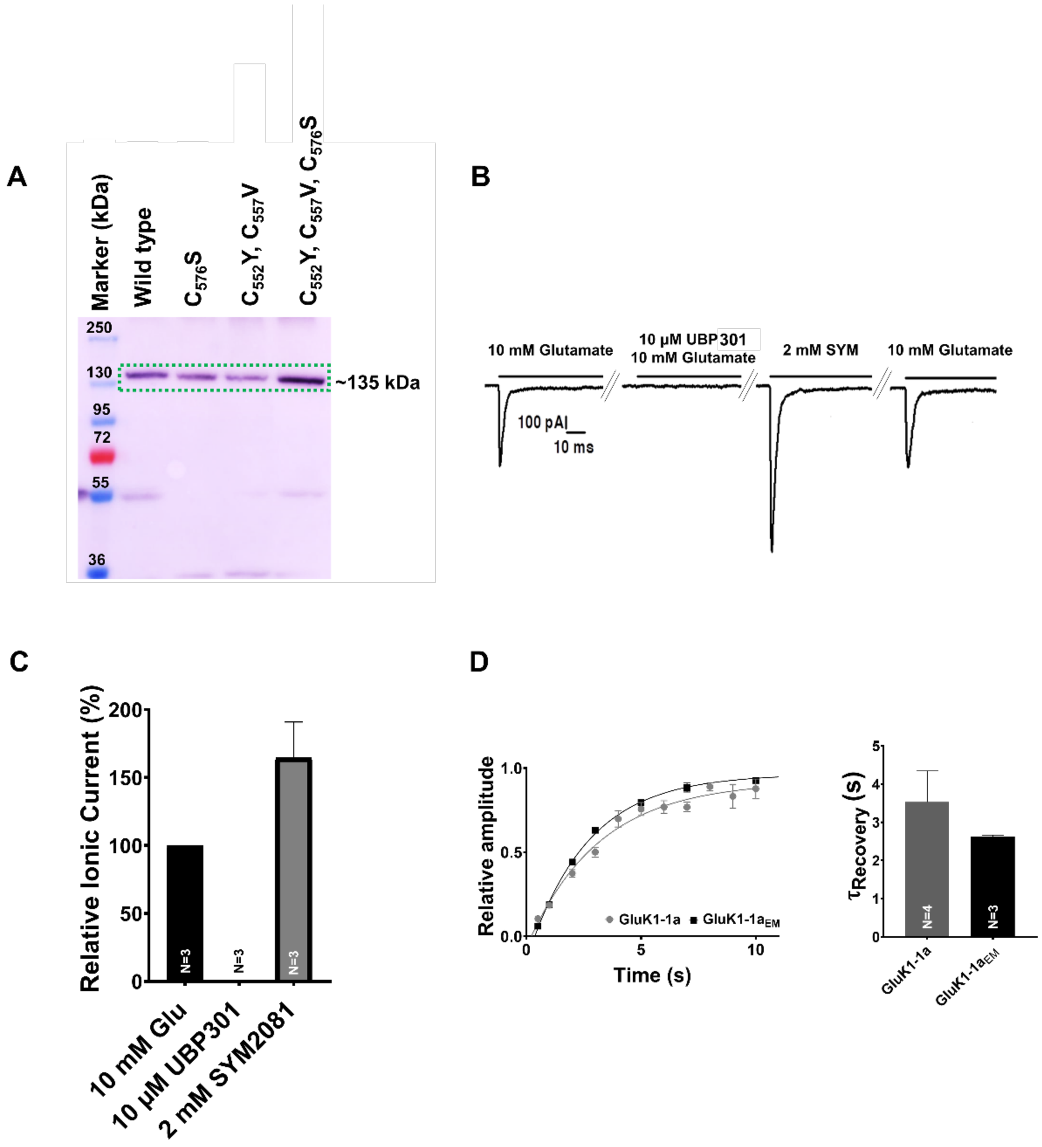


**Figure 6-figure supplement 3.** GluK1-1a construct optimization for structural studies and its gating properties. The optimized constructs were verified for expression. Here, three free cysteines mutations in the TM1 are represented as $C_{576}$S (1x Cys), $C_{552}$Y, $C_{557}$V (2x Cys), and $C_{552}$Y, $C_{557}$V, $C_{576}$S (3x Cys). 2x Cys ($C_{552}$Y, $C_{557}$V) mutant was used as GluK1-1a$_{EM}$ for structural studies. **(A)** Western blot probed with anti-His antibody (Cell Signaling Technology, USA) shows the expression of all the mutants with respect to the WT construct; Expected size of the polypeptide is indicated (MW: ~135 kDa). **(B)** Representative traces for whole-cell patch clamp recordings (36-48 h post-infection) HEK293 cells infected with GluK1-1a$_{EM}$ baculovirus is shown. The receptors showed activation by 10 mM Glutamate, blocked in the presence of the inhibitor 10 μM UBP301. Post-washing the reversible inhibitor, the receptor could undergo a similar activation and desensitization cycle in the presence of 2 mM SYM and 10 mM Glutamate. **(C)** Shows the graphical representation of the percentage of relative current in the presence of 10 μM UBP301, 2 mM SYM and 10 mM glutamate. **(D)** Electrophysiology profiles confirm that the GluK1-1a$_{EM}$ construct behaves similarly to the wild-type GluK1-1a in terms of recovery from desensitization. Error bars indicate mean ± SEM, N in each bar represents the number of cells used for analysis.

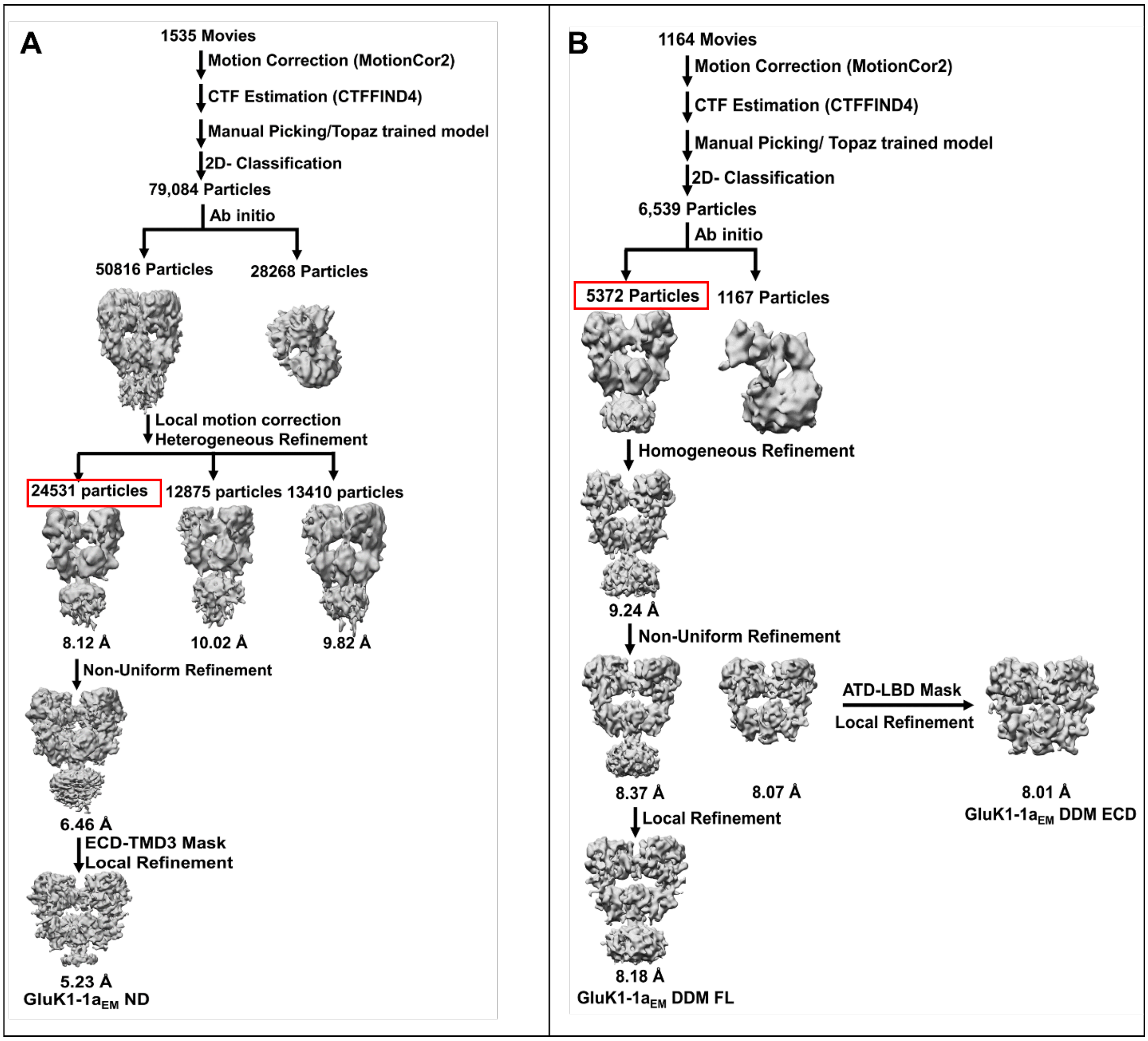


**Figure 6-figure supplement 4.** Single-particle cryo-EM data processing flow chart for GluK1-1a$_{EM}$ in nanodisc (ND) and detergent (DDM). **(A)** For the GluK1-1a$_{EM}$ ND dataset, 79,084 particles from the final 2D classification were used for initial 3D reconstruction into 2 classes to remove junk particles. Further, the good 50,816 particles were polished using local motion correction, and the initial 3D map was heterogeneously refined into 3 classes. The best map (24531 particles, highlighted in red box) was subsequently refined using non-uniform refinement followed by local refinement using the ECD-TMD3 mask to attain the final density map (GluK1-1a$_{EM}$ ND) resolution of 5.23 Å at 0.143 FSC. **(B)** 6,539 particles from the final 2D classification were used to determine *ab initio* 3D reconstruction of GluK1-1a$_{EM}$ DDM in

2 classes to remove broken particles. The 3D map was refined using good particles (5372, highlighted in red box) with homogenous refinement to obtain a resolution of 9.2 Å. Further, non-uniform refinement followed by local refinement was performed using full-length and extracellular domain (ECD=ATD+LBD) masks to get final resolutions of 8.2 Å (GluK1-1a$_{EM}$ DDM FL) and 8 Å (GluK1-1a$_{EM}$ DDM ECD) respectively.

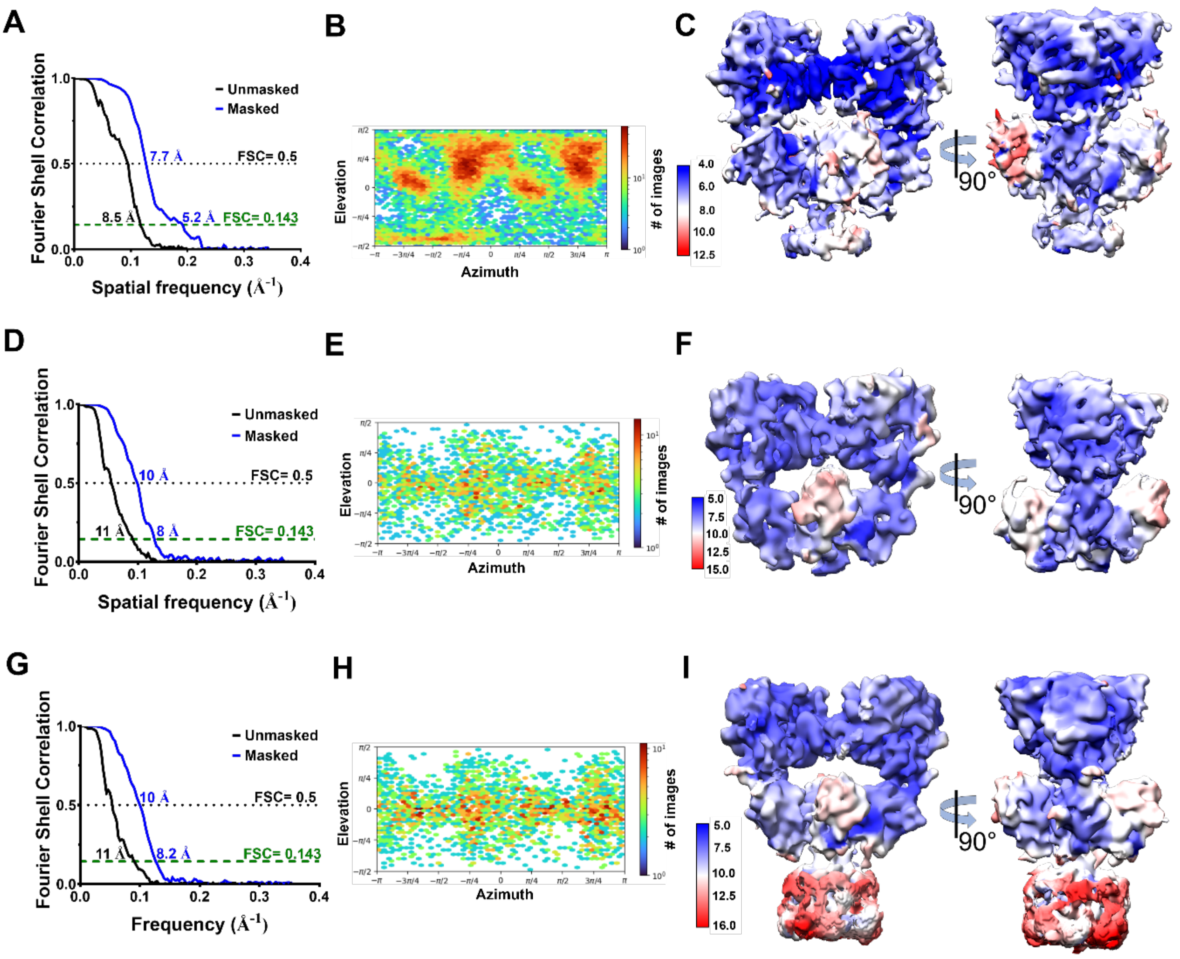


**Figure 6-figure supplement 5.** Estimation of resolution and particle distribution for GluK1-1a$_{EM}$ structures. **(A, D & G)** Show Fourier Shell Correlation curves at 0.143 and 0.5 cut-offs for GluK1-1a$_{EM}$ ND, GluK1-1a$_{EM}$ DDM ECD, and GluK1-1a$_{EM}$ DDM FL reconstructions, respectively, for unmasked (black) and masked (blue) maps estimated in cryoSPARCv3.1. **(B, E & H)** Show the angular distribution of particles for GluK1-1a$_{EM}$ ND, GluK1-1a$_{EM}$ DDM ECD, and GluK1-1a$_{EM}$ DDM FL maps, respectively, as produced by cryoSPARCv3.1. **(C, F & I)** Show the local resolution estimates for GluK1-1a$_{EM}$ ND (4-12.5 Å), GluK1-1a$_{EM}$ DDM ECD (5-15 Å), and GluK1-1a$_{EM}$ DDM FL (5-16 Å) maps respectively.

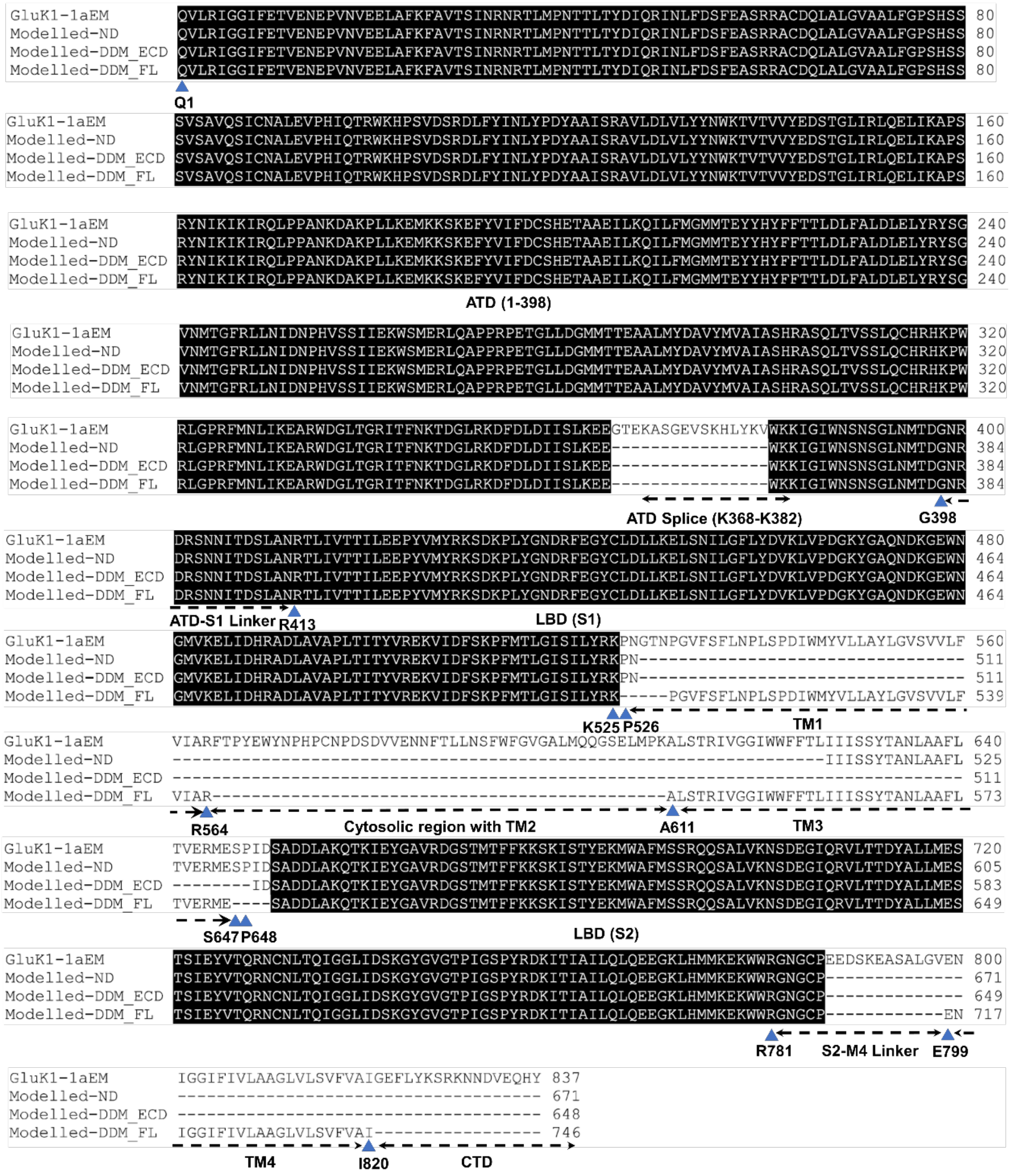

**Figure 6-figure supplement 6.** Sequence alignment for the three models presented in the study. Polypeptide chains modeled in the EM maps vs. the full GluK1-1a$_{EM}$ construct are shown. The beginning and ending residues of each modeled domain are indicated by blue triangles for GluK1-1a$_{EM}$ ND, GluK1-1a$_{EM}$ DDM ECD, and FL models. Missing residues are shown as dashed lines that could not be built due to resolution limitations.

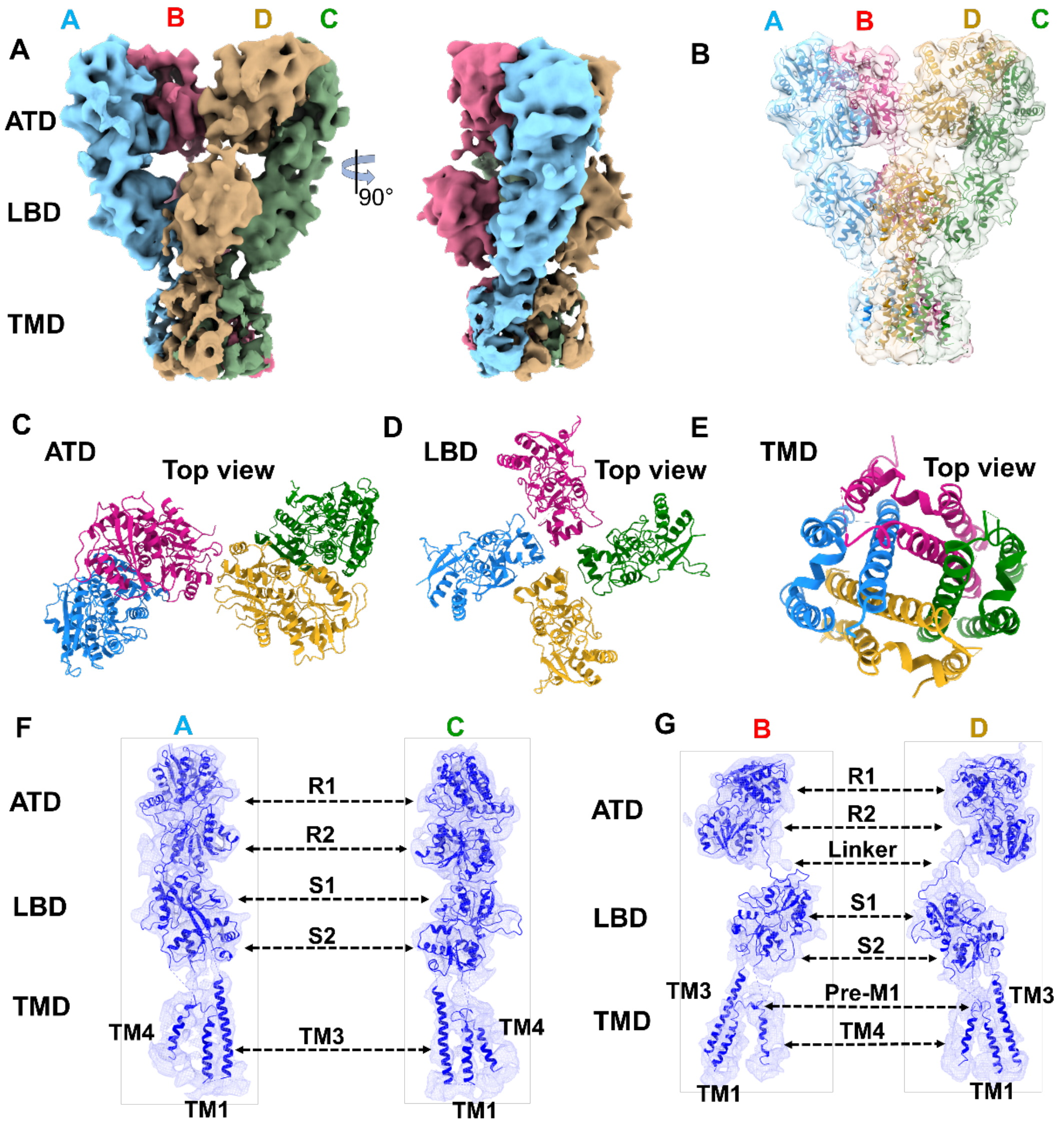


**Figure 6-figure supplement 7.** Cryo-EM map and model for GluK1-1a$_{EM}$ DDM FL-SYM complex. **(A)** Shows the front and side views of the segmented density map for the DDM solubilized GluK1-1a colored uniquely according to different chains. **(B)** Shows the atomic model fitted in the EM map. **(C, D & E)** Show the top views of ATD, LBD, and TMD layers. **(F & G)** Show the segmented map with fitted chains for each subunit, respectively. Sub-domains and helices of the TMD region are labelled.

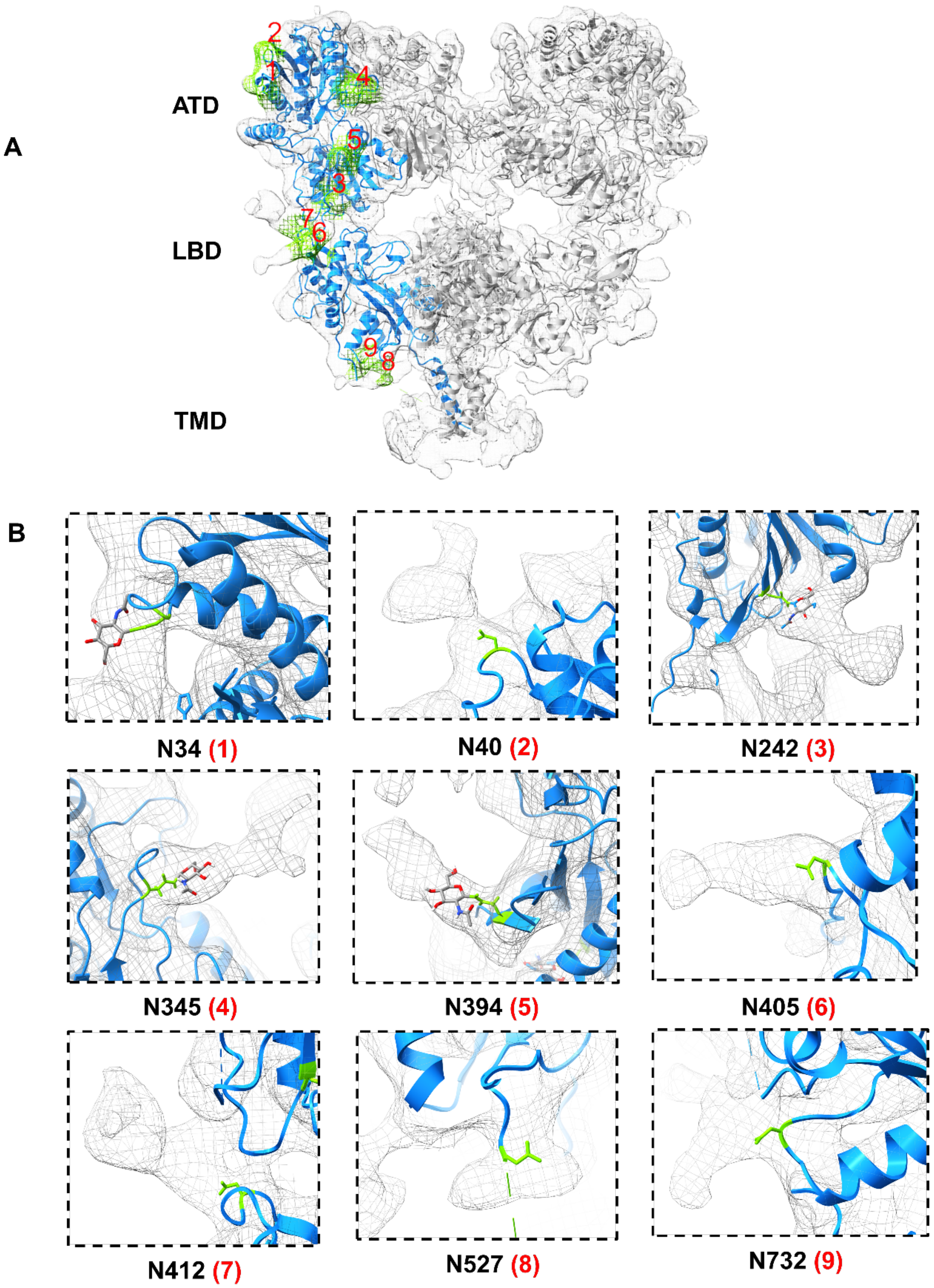


**Figure 6-figure supplement 8.** EM density map labelled to show predicted N-linked glycosylation sites (NXT) for GluK1-1a$_{EM}$. **(A)** Shows GluK1-1a$_{EM}$ ND with fitted model in EM density. Chain A is emphasized in blue, and the residual N-linked glycan densities are shown in green color with respective Asn residues labeled from 1-9. **(B)** Shows zoomed view of individual Asn residues with side chain shown as a stick model and the corresponding glycan density in mesh form. For $N_{34}$, $N_{242}$, $N_{345,}$ and $N_{394}$ residues, glycan density (NAG) observed in the GluK1-1a ATD crystal structure has been depicted with a conventional color scheme (C-gray, O- red, N- blue, H- white).

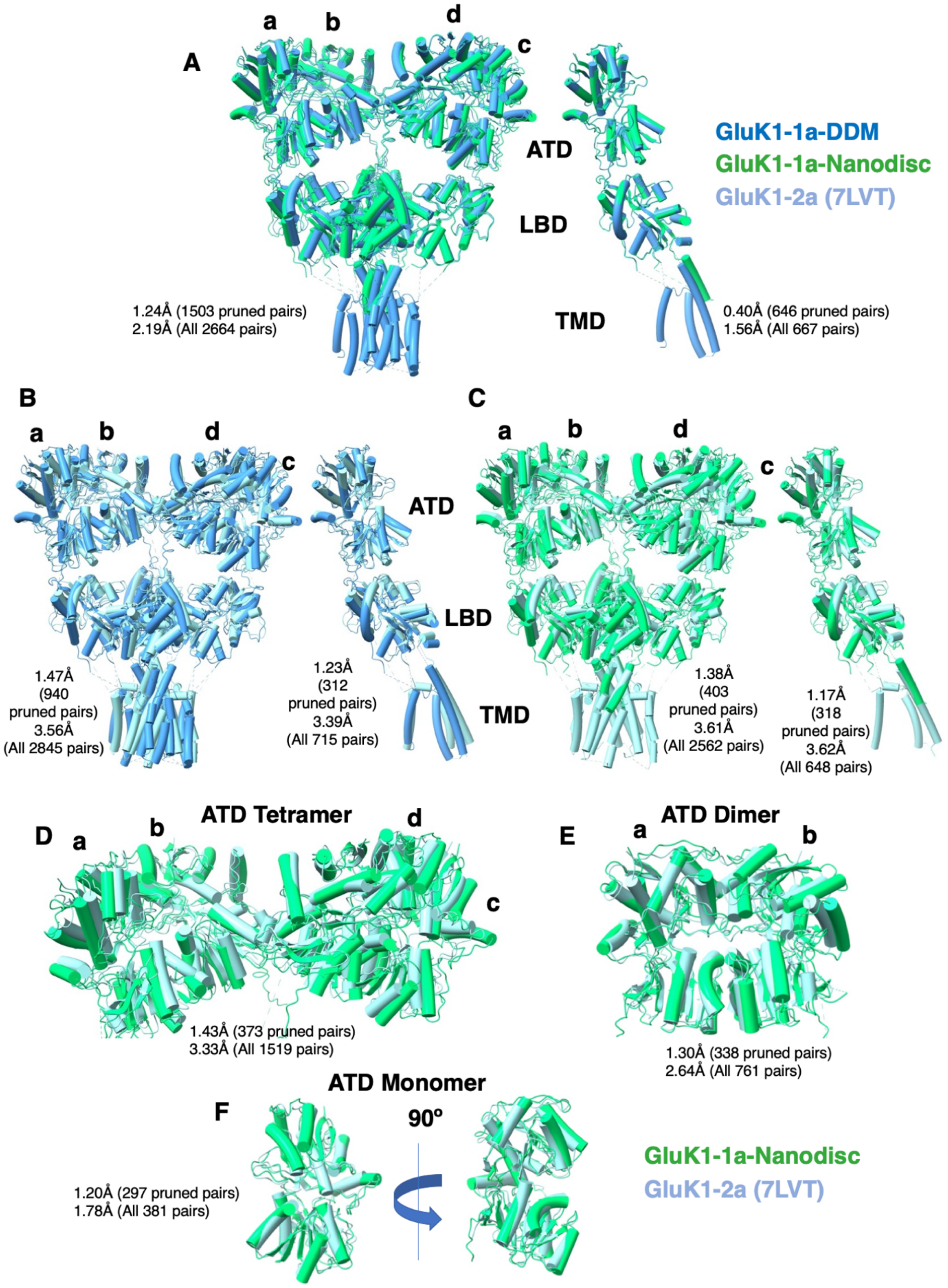


**Figure 6-figure supplement 9.** Comparison between GluK1-1a$_{EM}$ (detergent-solubilized or reconstituted in nanodiscs) and GluK1-2a (PDB-7LVT) in the desensitized state. Each panel

(A-F) illustrates a pairwise comparison with superimposed structures, where the root-mean-square deviation (RMSD) values, measured in Å, are indicated adjacent to each comparison. The structural comparison was carried out in ChimeraX and indicates significant structural similarities between all the protein models. The superimposition does not show significant differences in the arrangement at both ATD and LBD layers of GluK1-1a with respect to GluK1-2a.

| | GluK1-1a- 2S,4R-4-methyl glutamate | | |
|---|---|---|---|
| | **GluK1-1a$_{EM}$ ND** | **GluK1-1a$_{EM}$ DDM ECD** | **GluK1-1a$_{EM}$ DDM FL** |
| **Data Collection and Processing** | | | |
| **Microscope** | Titan Krios | Titan Krios | |
| **Voltage (keV)** | 300 | 300 | |
| **Number of micrographs** | 1535 | 1100 | |
| **Camera** | K2 | Falcon3 | |
| **Mode of recording** | Super resolution with energy filter (20eV slit) | Counting | |
| **Exposure time (s)** | 12 | 60 | |
| **Total dose ($e^-$/Å$^2$)** | 40.8 | 19.5 | |
| **Defocus range (µm)** | 1.8-3.2 | 2.0-3.2 | |
| **Pixel size (Å)** | 1.41 | 1.38 | |
| **Symmetry** | C1 | C1 | |
| **Initial particle number** | 1,97,908 | 13,750 | |
| **Final particle number** | 24531 | 5372 | |
| **Map resolution (Å)**<br>**FSC threshold** | 5.23<br>0.143 | 8.01<br>0.143 | 8.2<br>0.143 |
| **Refinement (Phenix)** | | | |
| **Initial model used (PDB code)** | (ATD), 3C32 (LBD), 5KUF(TM3) | (ATD), 3C32 (LBD) | (ATD), 3C32 (LBD), 5KUF (TMD) |
| **Model resolution (Å)**<br>**FSC threshold**<br>**Map-to model fit, CC_mask** | 5.1/7.3<br>0.143/0.5<br>0.71 | 7.2/9.0<br>0.143/0.5<br>0.69 | 7.8/9.1<br>0.143/0.5<br>0.73 |
| **Model composition** | | | |
| **Non-hydrogen atoms**<br>**Protein residues** | 21556<br>2684 | 20880<br>2596 | 23616<br>2948 |
| **R.m.s. deviations** | | | |
| **Bond lengths (Å)**<br>**Bond angles (°)** | 0.004<br>0.861 | 0.003<br>0.796 | 0.004<br>0.805 |
| **Validation** | | | |
| **MolProbity score**<br>**Clashscore** | 1.92<br>15.03 | 2.03<br>19.06 | 2.12<br>20.96 |
| **Ramachandran plot** | | | |
| **Favored (%)**<br>**Allowed (%)**<br>**Disallowed (%)** | 96.39<br>3.53<br>0.08 | 96.27<br>3.65<br>0.08 | 95.59<br>4.38<br>0.03 |
| **Rotamer outliers (%)** | 0.30 | 0.17 | 0.12 |
| **Cß outliers (%)** | 0.04 | 0 | 0.04 |

**Table 1: Cryo-EM data collection, refinement and validation for GluK1-1a$_{EM}$.**

**Figure 6- table supplement 1.** Cryo-EM data collection, refinement, and validation for GluK1-1A$_{EM}$.
